# Dopamine and DBS accelerate the neural dynamics of volitional action in Parkinson’s disease

**DOI:** 10.1101/2023.10.30.564700

**Authors:** Richard M. Köhler, Thomas S. Binns, Timon Merk, Guanyu Zhu, Zixiao Yin, Baotian Zhao, Meera Chikermane, Jonathan Vanhoecke, Johannes L. Busch, Jeroen G.V. Habets, Katharina Faust, Gerd-Helge Schneider, Alessia Cavallo, Stefan Haufe, Jianguo Zhang, Andrea A. Kühn, John-Dylan Haynes, Wolf-Julian Neumann

## Abstract

The ability to initiate volitional action is fundamental to human behaviour. Loss of dopaminergic neurons in Parkinson’s disease is associated with impaired action initiation, also termed akinesia. Both dopamine and subthalamic deep brain stimulation (DBS) can alleviate akinesia, but the underlying mechanisms are unknown. An important question is whether dopamine and DBS facilitate de novo build-up of neural dynamics for motor execution or accelerate existing cortical movement initiation signals through shared modulatory circuit effects. Answering these questions can provide the foundation for new closed-loop neurotherapies with adaptive DBS, but the objectification of neural processing delays prior to performance of volitional action remains a significant challenge.

To overcome this challenge, we studied readiness potentials and trained brain signal decoders on invasive neurophysiology signals in 25 DBS patients (12 female) with Parkinson’s disease during performance of self-initiated movements. Combined sensorimotor cortex electrocorticography (ECoG) and subthalamic local field potential (LFP) recordings were performed OFF therapy (*N*=22), ON dopaminergic medication (*N*=18) and ON subthalamic deep brain stimulation (*N*=8). This allowed us to compare their therapeutic effects on neural latencies between the earliest cortical representation of movement intention as decoded by linear discriminant analysis classifiers and onset of muscle activation recorded with electromyography (EMG).

In the hypodopaminergic OFF state, we observed long latencies between motor intention and motor execution for readiness potentials and machine learning classifications. Both, dopamine and DBS significantly shortened these latencies, hinting towards a shared therapeutic mechanism for alleviation of akinesia. To investigate this further, we analysed directional cortico-subthalamic oscillatory communication with multivariate granger causality. Strikingly, we found that both therapies independently shifted cortico-subthalamic oscillatory information flow from antikinetic beta (13-35 Hz) to prokinetic theta (4-10 Hz) rhythms, which was correlated with latencies in motor execution.

Our study reveals a shared brain network modulation pattern of dopamine and DBS that may underlie the acceleration of neural dynamics for augmentation of movement initiation in Parkinson’s disease. Instead of producing or increasing preparatory brain signals, both therapies modulate oscillatory communication. These insights provide a link between the pathophysiology of akinesia and its’ therapeutic alleviation with oscillatory network changes in other non-motor and motor domains, e.g. related to hyperkinesia or effort and reward perception. In the future, our study may inspire the development of clinical brain computer interfaces based on brain signal decoders to provide temporally precise support for action initiation in patients with brain disorders.

## Introduction

The conscious intent to initiate movement represents the core of our ability to exert autonomous control over our actions^1^. The underlying interplay of neural circuits builds the foundation for our capacity to interact with the world and adapt to our surroundings. The “‘decision’ to act” robustly elicits brain signals that precede motor output,^2^ which are observable from the single unit level^3^ to whole-brain activity with functional MRI.^4^ Nevertheless, the underlying neural dynamics that govern volitional action remain largely unknown.

Rodent studies employing optogenetics and fibre photometry have suggested that dopamine plays a pivotal role in volition and self-initiated movement.^5^ Dopaminergic neurons of the substantia nigra pars compacta (SNc) innervating the striatum increase their activity prior to spontaneous movement,^6^ while direct optogenetic activation of these neurons can increase the probability for movement initiation, suggestive of a causal role for dopamine in action initiation.

Parkinson’s disease is associated with a loss of dopaminergic neurons in the SNc that results in impaired movement initiation, also termed akinesia.^7^ Dopaminergic medication (levodopa) and subthalamic nucleus (STN) deep brain stimulation (DBS) can alleviate akinesia in Parkinson’s disease.^8^ Invasive neurophysiological recordings through DBS electrodes revealed that both dopamine and subthalamic DBS can shift akinetic pathological cortex–basal ganglia circuit synchrony to prokinetic states,^9^ but the shared mechanism of dopamine and DBS for action initiation is unclear.

In the present study, we leveraged multielectrode invasive neurophysiology recordings of preparatory brain signals combined with machine learning based brain signal decoding to investigate effects of dopamine and STN-DBS on the neural representation of motor intention in Parkinson’s disease. We provide physiological evidence for the importance of dopamine for volitional action initiation in humans and show that both dopaminergic medication and STN-DBS can shift oscillatory drive from cortex to the STN to theta (4-10 Hz) rhythms, providing mechanistic insight into cortex–basal ganglia interactions that govern and impede action initiation in Parkinson’s disease.

## Materials and Methods

### Experimental model and subject details

25 patients (12 female) diagnosed with idiopathic Parkinson’s disease of primary akinetic-rigid motor phenotype with clinical indication for deep brain stimulation (DBS) were enrolled (Table 1). Patients were enrolled at the Department of Neurosurgery of the Beijing Tiantan Hospital, Capital Medical University (*N*=9) or at the Department of Neurosurgery at Charité – Universitätsmedizin Berlin (*N*=16). The mean age of subjects was 62.4±8.4 years and mean disease duration was 9.9±4.2 years. DBS leads were implanted into the subthalamic nucleus (STN) of both hemispheres in all patients. A subdural electrocorticography (ECoG) electrode was implanted unilaterally on the hand knob area of the primary motor cortex of all patients for research purposes.

**Table 1:** Participant information. DBS: deep brain stimulation; F: female; M: male; DD: disease duration in years; LEDD: levodopa-equivalent daily dose in mg; DBS: deep brain stimulation; ECoG: electrocorticography; MT: Medtronic; BS: Boston Scientific; PINS: PINS Medical; AT: Ad-Tech; SMT: Sinovation Medical Technologies

| ID | Age | Sex | DD<br>[y] | LEDD<br>[mg] | DBS lead model | ECoG model | ECoG<br>hemisphere |
| --- | --- | --- | --- | --- | --- | --- | --- |
| EL002 | 62 | M | 10 | 1032 | BS Vercise Cartesia | AT TS06R-AP10X-0W6 | left |
| EL003 | 56 | F | 15 | 600 | MT 3389 | AT TS06R-AP10X-0W6 | left |
| EL004 | 45 | F | 2 | 1462 | BS Vercise Cartesia | AT TS06R-AP10X-0W6 | left |
| EL005 | 43 | F | 4 | 1221 | MT SenSight Short | AT TS06R-AP10X-0W6 | right |
| EL006 | 57 | M | 6 | 1000 | MT SenSight Short | AT TS06R-AP10X-0W6 | right |
| EL007 | 55 | M | 3 | 850 | MT SenSight Short | AT DS12A-SP10X-000 | right |
| EL008 | 66 | M | 8 | 858 | BS Vercise Cartesia X | AT TS06R-AP10X-0W6 | left |
| EL009 | 59 | M | 7 | 1600 | MT SenSight Short | AT TS06R-AP10X-0W6 | left |
| EL010 | 66 | F | 12 | 900 | MT SenSight Short | AT TS06R-AP10X-0W6 | right |
| EL011 | 67 | M | 7 | 949 | MT SenSight Short | AT TS06R-AP10X-0W6 | right |
| EL012 | 54 | M | 12 | 866 | MT SenSight Short | AT TS06R-AP10X-0W6 | right |
| EL013 | 64 | M | 9 | 800 | MT SenSight Short | AT DS12A-SP10X-000 | right |
| EL014 | 52 | M | 14 | 1750 | MT SenSight Short | AT TS06R-AP10X-0W6 | right |
| EL015 | 60 | M | 14 | 708 | MT SenSight Short | AT TS06R-AP10X-0W6 | right |
| EL016 | 73 | M | 20 | 1630 | MT SenSight Short | AT TS06R-AP10X-0W6 | right |
| EL017 | 65 | F | 8 | 1150 | MT SenSight Short | AT TS06R-AP10X-0W6 | right |
| FOG006 | 66 | F | 10 | 513 | PINS L301 | SMT PSE-8A | right |
| FOG008 | 70 | M | 12 | 688 | PINS L301 | SMT PSE-8A | right |
| FOG010 | 73 | F | 9 | 1439 | PINS L301 | SMT PSE-8A | right |
| FOG012 | 59 | F | 9 | 700 | PINS L301 | SMT PSE-8A | right |
| FOG014 | 76 | M | 8 | 1351 | PINS L301 | SMT PSE-8A | right |
| FOG016 | 66 | F | 15 | 669 | PINS L301 | SMT PSE-8A | right |
| FOG021 | 66 | F | 15 | 925 | PINS L301 | SMT PSE-8A | right |
| FOG022 | 67 | F | 9 | 1000 | PINS L301 | SMT PSE-8A | right |
| FOGC001 | 72 | F | 10 | 675 | PINS L301 | SMT PSE-8A | right |

### Ethics declaration

The research and brain signal recordings presented in this manuscript were performed according to the standards set by the declaration of Helsinki and after approval of the local independent ethics review board at the included academic centres (Berlin and Beijing). All patients provided informed consent to participate in the respective research according to the declaration of Helsinki. For data from Berlin, the studies were approved by the ethics committee at Charité – Universitätsmedizin Berlin (EA2/129/17). The data was collected, stored, and processed in compliance with the General Privacy Protection Regulation of the European Union. The data collection for Beijing was approved by the independent review board of Beijing Tiantan Hospital (KY 2018-008-01), registered in the Chinese Clinical Trial Registry (ChiCTR1900026601) and conducted under the supervision of an authoritative third party (China National Clinical Research Center for Neurological Diseases).

### DBS and ECoG placement

DBS implantation followed a 2-step approach: In a first surgery, DBS leads were placed stereotactically after co-registering pre-operative MRI and CT images. A single ECoG electrode was placed subdurally onto one hemisphere after minimal enlargement (∼2 mm) of the frontal burr hole. The ECoG electrode was aimed posteriorly toward the hand knob region of the motor cortex. The ECoG electrode was placed either on the right hemisphere (Beijing; right=9) or on the hemisphere ipsilateral to the implantable pulse generator (Berlin; right=11, left=5). All electrodes were then externalized through the burr holes via dedicated externalization cables. Patients remained on the ward for a duration of 4-7 days up until the second surgery. During the time between surgeries, electrophysiological recordings were performed. In the second intervention, externalization cables of the DBS leads were replaced with permanent cables that were tunnelled subcutaneously and connected to a subclavicular implantable pulse generator. ECoG electrodes were removed via the burr hole during the second surgery.

### Anatomical localization of electrodes

DBS and ECoG electrodes were localized using standard settings in Lead-DBS.^10^ In brief, pre-operative MRI and post-operative CT images were co-registered, corrected for brain shift, and normalized to MNI space (Montreal Neurological Institute; MNI 2009b NLIN ASYM atlas).

Electrode artifacts were marked manually and MNI coordinates were extracted. See Supplementary Fig. 1 for reconstructed electrode localizations. Individual electrode contacts were then assigned to one of either parietal, sensory, motor, or prefrontal cortex regions based on proximity to these regions using the AAL3 parcellation.^11^ Equally, bipolar ECoG channels were assigned to one of parietal, sensory, motor, or prefrontal cortex based on proximity of the mean Euclidian distance between the MNI coordinates of the two used electrode contacts to the respective cortical regions.

### Experimental paradigm

Study participants performed self-initiated movements with the upper limb contralateral to the ECoG hemisphere. 16 subjects performed a wrist rotation on a custom-built “rotameter” device that translated the degree of rotation to voltage (see Figure 1D for schematic). 9 subjects performed a button press with their left index finger that elicited a transistor-transistor logic (TTL) signal. Participants were instructed to pause for about 10 seconds between movements, but to avoid counting seconds in between. Subjects performed an average of 66±36.25 movements.

**Figure 1.**
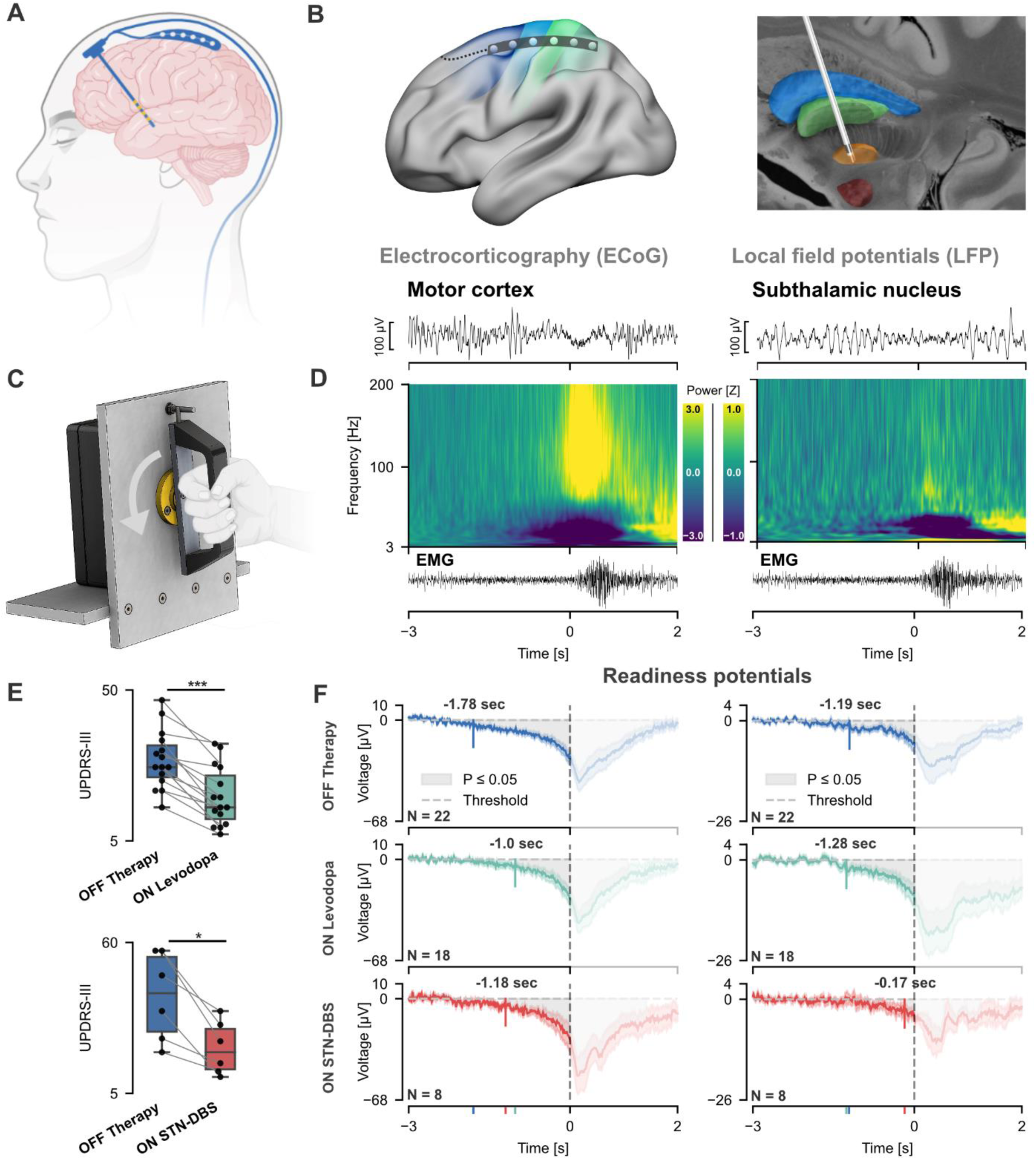
Sensorimotor electrocorticography and subthalamic local field potentials during self-initiated movements in Parkinson’s disease patients. **(A)** 25 Parkinson’s disease patients were implanted with bilateral deep brain stimulation (DBS) leads and a single electrocorticography (ECoG) strip. **(B)** ECoG strips were placed on the sensorimotor cortex and deep brain stimulation (DBS) electrodes placed into the dorsolateral part of the subthalamic nucleus. Exemplar traces show ECoG activity from the motor cortex and subthalamic local field potentials (STN-LFP) from the hemisphere contralateral to movement during a single movement. **(C)** Schematic of the rotational handle used by Parkinson’s disease patients in the Berlin cohort. **(D)** Oscillatory activity from the hemisphere contralateral to movement recorded OFF therapy, averaged across patients (*N*=22) and EMG of the brachioradial muscle during a self-initiated movement. **(E)** UPDRS-III (Unified Parkinson’s Disease Rating Scale) scores during experimental sessions of patients recorded both OFF and ON levodopa (29/18±8/8, *P*=6×10-5, *N*=15; Supplementary Table 1) and UPDRS-III scores at 12 months post-implantation for patients recorded both OFF therapy and ON subthalamic deep brain stimulation (STN-DBS; 41/22±14/8, *N*=7 – unavailable in *N*=1; Table S2). **(F)** Readiness potentials of motor cortex (left) and STN-LFP signals (right) contralateral to movement side, averaged across patients. Motor cortex readiness potentials differed significantly from baseline between −1.78 and 1.55 s (OFF therapy), −1.0 and 1.09 s (ON levodopa), and −1.18 and 1.51 s (ON STN-DBS). Subthalamic readiness potentials differed significantly from baseline between −1.19 and 1.32 s (OFF therapy), −1.28 and 2.0 s (ON levodopa), and −0.17 and 1.2s (ON STN-DBS; all *P*≤0.05, cluster corrected). Data are represented as mean ± SEM.

Participants performed the experimental task either under their current clinical intake of dopaminergic medication (ON levodopa; see Table 1 for details of levodopa-equivalent daily doses at time of recording) or after at least 12 hours of withdrawal of all dopaminergic medication (OFF therapy). A total of 22 subjects performed the task OFF therapy and 18 subjects ON levodopa. 16 subjects performed the task both OFF therapy and ON levodopa. An additional eight subjects that performed the task OFF therapy also performed the task after withdrawal of medication but under high-frequency DBS of the STN (ON STN-DBS). UPDRS-III scores at 12 months post-implantation for patients recorded both OFF therapy and ON STN-DBS were 41/22±14/8 (Figure 1H; *N*=7 – unavailable in *N*=1; see Table S2 for details). Contacts and stimulation parameters to be used during recording were determined in a monopolar clinical review. Clinically effective contacts were chosen while avoiding stimulation-induced side-effects altogether. DBS was applied at 130 Hz with 60 µs pulse-width, with a mean amplitude per subject of 1.88±0.55 mA (see Table S2 for details). 6 participants received bilateral monopolar stimulation. Due to side effects, 2 participants received unilateral DBS in the hemisphere contralateral to the extremity used in the task, one receiving monopolar and one bipolar stimulation.

### Electrophysiology recordings

All ECoG data and subthalamic local field potentials (STN-LFP) were recorded in the days between first and second surgical intervention. Subdural ECoG strips had either 6 (*N*=14), 8 (*N*=9), or 12 contacts (*N*=2). STN-LFP were recorded from one of 4 DBS lead models implanted with either 4 (*N*=10), 8 (*N*=14) or 16 (*N*=1) contacts (Table 1).

All data were amplified and digitized with either a NeuroOmega (Alpha Omega Engineering; 1376 Hz sampling rate), Saga64+ (Twente Medical Systems International; 4000 Hz sampling rate) or Neurofax EEG-1200 (Nihon Kohden; 2000 Hz sampling rate) device. ECoG signals and subthalamic local field potentials (STN-LFP) were hardware-referenced to the lowermost contact of the DBS electrode (ipsilateral to movement). In a small number of cases where excessive noise was visible before recording, either a different STN contact or an average of 2 ECoG contacts were used as hardware reference. Additionally, accelerometry, bipolar EMG, and TTL (button press) or rotameter (wrist rotation) signals were recorded. Data was saved to disk for offline processing and converted to iEEG-BIDS^12^ format. Offline processing was, unless noted otherwise, performed with custom python scripts using the MNE-Python,^13^ MNE-BIDS,^14^ NumPy,^15^ Pandas,^16^ SciPy,^17^ toolboxes and MATLAB scripts using the FieldTrip^18^ and SPM12 (https://www.fil.ion.ucl.ac.uk/spm/software/spm12/) toolboxes.

### EMG annotation

EMG was recorded from brachioradialis for wrist rotations and from finger flexors for button presses. The root mean square (RMS) of EMG was calculated using a sliding window of 100 ms length centred around each sample. EMG onset and end for each trial were manually annotated by inspection of the RMS data. In 3 subjects, noise in EMG was too high for adequate interpretation and in 1 subject, no EMG was recorded. Onset and end of the rotation and button press (TTL) signal were manually annotated for all available recordings. The mean difference between EMG and rotation/button press onset, and EMG and rotation/button press end, were calculated in recordings where both data were present. For recordings where no adequate EMG signal was available, EMG onset and end were estimated by subtracting the mean difference from all other recordings of the respective group (rotation/button press).

### Artifact rejection

ECoG and STN-LFP data were high-pass filtered at 3 Hz and epoched between −3 and 2 s relative to EMG onset. All trials were visually inspected and trials containing significant high-amplitude artifacts, for example due to cable movement during motor activity, were marked and excluded from further analysis.

### Readiness potentials

ECoG contacts located on the motor cortex and STN-LFP signals were referenced to the lowermost DBS contact of the electrode ipsilateral to the moved hand if this had not been the hardware reference during recording. Raw signals were filtered between 0.1 and 40 Hz and epochs were extracted from −3 to 2 s relative to EMG onset for each movement trial. The signals of each channel and epoch individually were baseline corrected by subtracting the mean of the baseline period (−3 to −2 s relative to EMG onset). Epochs were then averaged across channels for each subject and ECoG/STN-LFP channels separately.

### Time-frequency analysis

Raw data was re-sampled to 500 Hz, referenced in a bipolar montage, and notch-filtered to remove line noise (50 Hz and all upper harmonics). For ECoG data, bipolar channels located on motor cortex were selected (see section Anatomical localization of electrodes for details). Epochs from −3 to 2 s relative to EMG onset were extracted. Power was calculated from epochs using Morlet wavelets (3-200 Hz, 1 Hz steps, 7 cycles). Power was averaged across trials within each subject and baseline correction was applied to each channel and epoch individually by subtracting the mean and dividing by the standard deviation of the baseline.

### Classification of motor intention

We processed data in a real-time compatible fashion using the open-source toolbox py_neuromodulation.^19^ In 100 ms steps, data batches of the previous 500 ms were processed. ECoG and STN-LFP signals were referenced in a bipolar setup and notch-filtered to remove line noise (50, 100 and 150 Hz). Band power was calculated for each bipolar channel via fast Fourier transform with a frequency resolution of 1 Hz and power at each frequency was normalized to samples from the previous 10 s. Band power at each time step was then averaged for 7 frequency bands: theta (4-7 Hz), alpha (8-12 Hz), low beta (13-20 Hz), high beta (21-35 Hz), low gamma (60-80 Hz), high gamma (90-200 Hz), high frequency activity (201-400 Hz).

To ensure that the target activity pattern has relevant modulation, while excluding activity after movement onset, we defined the single sample 100 ms prior to EMG onset as our motor intention decoding target. Samples between −3 and −2 s prior to EMG onset were defined as rest for model training (Figure 2A). Linear discriminant analysis (LDA) classifiers (scikit-learn^20^ implementation with least squares solver, automated shrinkage with equal class prior probabilities) were trained to separate motor intention from rest samples for each subject individually and on features across all available recording channels from ECoG and STN-LFP signals separately. For example, a classifier trained on ECoG data of a single recording with 5 bipolar ECoG channels would use 35 features at each time sample (5 channels x 7 frequency bands). Additionally, LDA classifiers were trained for each individual ECoG channel separately. As an example, a classifier trained in this analysis would use 7 features for classification (1 bipolar channel x 7 frequency bands). To avoid information leakage from training to testing data, classifiers were trained in a leave-one-group-out approach for each recording, where each separate movement trial represented a separate group. For example, if a participant had performed 50 movement trials in a single recording, the classifier was trained and tested in a 50-fold group cross-validation. In each fold, all samples from a single trial were left out and the classifier was trained on the remaining 49 trials. Then, classifications for the samples of the left-out trial were obtained from the trained classifier and balanced accuracy was calculated as metric of performance. The balanced accuracy was then averaged across all folds to obtain a single metric for each recording.

**Figure 2.**
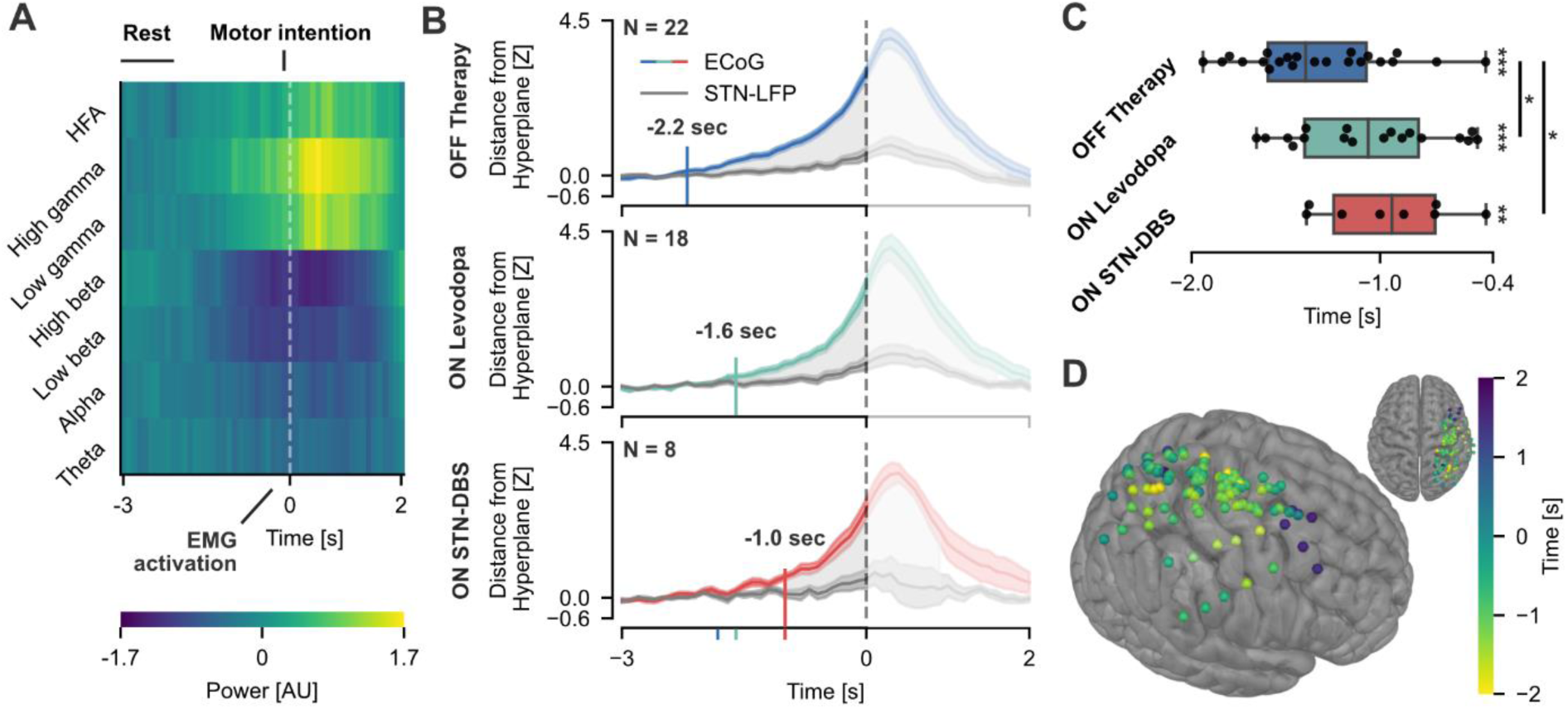
Dopamine and subthalamic deep brain stimulation reduce motor intention to execution delays in Parkinson’s disease. **(A)** Features of a single motor cortex channel averaged across trials. **(B)** Classifier outputs averaged across subjects. Classifier outputs of electrocorticography (ECoG) and subthalamic local field potentials (STN-LFP) differed between −2.2 to 1.7 s (OFF therapy), −1.6 to 1.6 s (ON levodopa) and −1.0 to 0.9 s (ON subthalamic deep brain stimulation [STN-DBS]; all *P*≤0.05, cluster corrected). Data are represented as mean ± SEM. **(C)** Time of motor intention of single subjects derived from ECoG classifier outputs. **(D)** Time of motor intention derived from *single-channel* ECoG classifier outputs. Left hemispheric channels were flipped onto the right hemisphere. \**P*≤0.05; \*\**P*≤0.01; \*\*\**P*≤0.001

Additionally, for each left-out trial the distance from the hyperplane separating rest and motor intention classes was calculated for samples between −3 s prior to 2 s after EMG onset as a proxy for the certainty of motor intention classification. Data were then normalized by subtracting the mean and dividing by the standard deviation of the baseline period (−3 s to −2 s) and average across trials within each recording. For each therapeutic condition (OFF therapy, ON levodopa, ON STN-DBS), the mean distance from hyperplane per recording at each time step between −3 s to 2 s was compared for ECoG and STN-LFP classifiers using two-sided permutation tests. P-values were corrected for multiple comparisons using cluster-based permutation testing (see section Statistical analysis for details).

To allow for group comparisons across therapeutic conditions, we determined the earliest significant deviation within subjects across baseline corrected trials (hereafter termed *time of motor intention*). For each recording, baseline corrected samples at each time step between −3 s to 2 s were tested against zero (x > 0) using a one-sided permutation test. P-values were corrected for multiple comparisons using cluster-based permutation testing (see section Statistical analysis for details). The time of the first statistically significant sample was then termed *time of motor intention*.

### Time-reversed Granger causality

Raw data was high-pass filtered at 1 Hz, re-sampled to 200 Hz, referenced in a bipolar montage, and notch-filtered to remove line noise (50 Hz). STN-LFP channels and bipolar ECoG channels located on motor cortex were selected (see section Anatomical localization of electrodes for details), all contralateral to the side of movement, and epochs from −3 to 2 s relative to EMG onset were extracted. Time-frequency estimates of multivariate^21^ time-reversed Granger causality^22^ (TRGC) were computed using a modified implementation of the MNE-Connectivity toolbox (https://mne.tools/mne-connectivity/stable/index.html). TRGC was calculated on time-frequency cross-spectral densities derived using Morlet wavelets (4 - 35 Hz, 1 Hz frequency resolution, 7 cycles) within the motor intention period (−2 - 0 s), and vector autoregressive models with orders of 30. Following the definition of TRGC in Winkler et al.^22^, Granger causality (GC) scores with motor cortex channels as seeds and STN channels as targets were obtained, and GC scores with STN channels as seeds and motor cortex channels as targets were subtracted to obtain net GC scores. Subsequently, the net GC scores from time-reversed data were obtained and subtracted from the original net GC scores to give TRGC scores. The final Granger scores therefore represent the net drive of information flow between motor cortex and STN, corrected for spurious estimates of connectivity arising from weak data asymmetries such as volume conduction and measurement noise.^22^ The connectivity values were smoothed within each recording with a 300 ms smoothing kernel and downsampled to 10 Hz for direct statistical comparison across OFF therapy, ON levodopa and ON STN-DBS cohorts. Figures related to Granger causality analyses were created using custom code in MATLAB R2022a.

### Statistical analysis

All statistical tests were performed using non-parametric permutation tests with at least 10,000 permutations and a value of α=0.05 for significance. P-values in time series and time-frequency series were corrected for multiple dependent comparisons using cluster-based permutation tests with at least 2,000 permutations.

### Code availability

All code is made publicly available under https://github.com/richardkoehler/paper-motor-intention.

## Results

### Subjects

Invasive neural population activity from sensorimotor electrocorticography (ECoG) and subthalamic deep brain stimulation (DBS) electrodes was recorded during self-initiated movements in 25 patients with Parkinson’s disease (Figure 1A-D). Patients were recruited from two specialized DBS centres at Charité – Universitätsmedizin Berlin in Germany and Tiantan Hospital Beijing in China and performed either rotational hand movements (*N*=16; Berlin) or index finger button presses (*N*=9; Beijing). Patients performed 66±36.25 movements (mean ± standard deviation) at their individual pace without any external cues that were recorded with EMG. Experiments were performed after withdrawal (*N*=22; 12/10 male/female) or administration of levodopa (*N*=18; 11/7 male/female). Dopaminergic medication led to an acute and significant alleviation of parkinsonian motor signs (UPDRS-III OFF/ON levodopa 29/18±8/8, *P*=6×10^-5^, available in *N*=15; Figure 1E; Supplementary Table 1). In a subset of eight patients (Berlin), ECoG-LFP recordings were repeated during high-frequency DBS of the STN (OFF levodopa; 130 Hz, 60 µs, clinically effective contact, mean amplitude 1.88±0.55 mA; available in *N*=8; Table S2). ECoG and STN-LFP activity from the hemisphere contralateral to movement showed typical movement-related oscillatory changes with decreased alpha (∼8-12 Hz) and beta (∼13-35 Hz) activity and increased gamma (∼60-90 Hz) and broadband high frequency activity (∼90-200 Hz).

### Readiness potentials in motor cortex are shorter with dopamine and STN-DBS

To investigate pathophysiological changes related to Parkinson’s disease and to study the modulatory effect of dopaminergic medication and STN-DBS on the neural representation of movement initiation, we first extracted readiness potentials (also called local motor potential or Bereitschaftspotential) from motor cortex and STN activity contralateral to the moved hand. Across subjects, readiness potential amplitude in motor cortex was significantly above baseline relative to EMG onset and significantly stronger in amplitude than STN-LFP across all conditions (all *P*≤0.05; not shown). In the medication and stimulation OFF therapy condition (*N*=22) the earliest significant readiness potential deflections in motor cortex started on average as early as −1.78 s relative to EMG onset (Figure 1F; *P*≤0.05; cluster corrected).

We hypothesized that this could be indicative of a pathologically prolonged movement initiation period in the hypodopaminergic parkinsonian state and analysed the effect of levodopa and STN-DBS therapy. Indeed, the latency between readiness potential onset and motor execution in the ON levodopa (*N*=18) condition was 780 ms shorter with earliest significant signal deflections observable at −1.0 s relative to EMG onset. Similarly, the ON STN-DBS (*N*=8) condition showed a 600 ms shorter latency between readiness potential onset and motor execution with earliest deflections in the ON STN-DBS (OFF levodopa) condition occurring at −1.18 s (*N*=8; all *P*≤0.05, cluster corrected). Readiness potentials in the STN, despite being significantly lower in amplitude when compared to ECoG, were also found consistently above baseline at −1.19 s (OFF therapy; *N*=22), −1.28 s (ON levodopa; *N*=18) and −0.17 s (ON STN-DBS; *N*=8; all *P*≤0.05, cluster corrected; Figure 1F).

### ECoG oscillations predict motor output before movement execution

We aimed to get a more reliable estimate of the neural representation of motor intention in Parkinson’s disease and its modulation with dopamine and STN-DBS, and to confirm our results with an additional analysis that is independent of readiness potential. For this, we trained multivariate machine learning classifiers on the oscillatory premotor patterns from all available channels. ECoG and STN-LFP signals were decomposed into seven frequency bands (theta (4-7 Hz), alpha (8-12 Hz), low beta (13-20 Hz), high beta (21-35 Hz), low gamma (60-80 Hz), high gamma (90-200 Hz), high frequency activity (HFA; 201-400 Hz)) at a temporal resolution of 100 ms.

To ensure that the target activity pattern has relevant preparatory modulation, while excluding activity after movement onset, we defined activity 100 ms prior to EMG onset as our motor intention decoding target. Samples between −3 and −2 s prior to EMG onset were defined as rest for model training (Figure 2A). Linear discriminant analysis (LDA) classifiers were trained to separate motor intention from resting activity. The classifier output (distance from the hyperplane separating rest and motor intention classes) was then calculated for each sample on left out cross-validation folds between −3 s prior to 2 s after EMG onset for each trial as a proxy for the certainty of motor intention classification.

Significant deviations of classifier output across subjects preceded the longest readiness potentials and were significantly higher for ECoG than STN-LFP classifiers (Figure 2B). Significant clusters started earliest OFF therapy −2.2 s to 1.7 s (*N*=22), followed by ON levodopa −1.6 s to 1.6 s (*N*=18) and ON STN-DBS −1.0 s to 0.9 s (*N*=8; all *P*≤0.05, cluster corrected). Before investigating specific delays from earliest decoding time-points and comparison across cohorts, we ensured that classifier accuracy was satisfactory. Motor intention and rest features from ECoG were classified significantly above chance (50%) across subjects with a mean balanced accuracy of 79.7±7.4% (*P*=4×10^-6^), 77.2±9.9% (*P*=8×10^-6^) and 77.2±8.0% (*P*=8×10^-3^) in recordings OFF therapy, ON levodopa and ON STN-DBS respectively (Supplementary Fig. 2A). Classification accuracy did not differ across medication and stimulation conditions (Supplementary Fig. 2A-B; all *P*>0.05), but ECoG classifiers were significantly more accurate than STN-LFP classifiers in all therapeutic conditions (Supplementary Fig. 2C).

### Dopamine and STN-DBS shorten latencies from motor intention to execution

To allow for robust and direct group comparisons across therapeutic conditions, we determined the earliest significant deviation of classifier output at the single-subject level across baseline corrected trials (hereafter termed *time of motor intention*). Following this analysis, the mean *time of motor intention* derived from ECoG occurred significantly before EMG onset (Figure 2C) at −1.3±0.4 s (OFF therapy; *N*=22; *P*=4×10^-6^), −1.1±0.4 s (ON levodopa; *N*=18; *P*=8×10^-6^) and −1±0.3 s (ON STN-DBS; *N*=8; *P*=8×10^-3^). Mirroring the observed pattern of readiness potential onset, but now statistically comparable across therapeutic conditions, the mean time of motor intention in ECoG was earlier for recordings OFF therapy than ON levodopa (Δ=−270 ms, *P*=0.039) and OFF therapy than ON STN-DBS (Δ=−360 ms, *P*=0.03). The effect for OFF versus ON levodopa was reproduced when only subjects recorded in both conditions were considered (Supplementary Fig. 2D; *P*=0.046, ΔLevodopa−140±470 ms).

For STN-LFP recordings, mean *time of motor intention* was not reliably estimable before EMG onset across all subjects (Supplementary Fig. 2G) indicative of the conservative nature of this measure. We therefore did not consider the STN-LFP for further analysis of motor intention times.

In summary, our results provide evidence for a prolonged cortical representation of motor intention in the parkinsonian OFF state that may reflect the impairment in movement initiation. Both dopaminergic medication and STN-DBS may reduce motor intention to movement execution delays, potentially reflective of their ability to alleviate akinesia in Parkinson’s disease.

### Therapeutic modulation of motor intention signals is regionally specific to motor cortex

To investigate anatomical specificity, we calculated the *time of motor intention* (as above) for every channel separately (Figure 2D; OFF therapy: *N*=120 channels). Channels were assigned to one of either parietal, sensory, or motor cortex based on proximity using the Automated Anatomical Label (AAL3) parcellation.^11^ Due to the location of the burr hole in STN-DBS surgery, only few ECoG channels (*N*=13 from 11 patients) were located anterior to motor cortex (mostly middle frontal gyrus), which is prefrontal areas could not be included in further analyses. *Time of motor intention* occurred significantly before EMG onset for all three parietal, sensory and motor cortical regions (all *P*≤1×10^-3^) but was significantly earlier in motor cortex than in parietal (Δ=−520 ms, *P*=5×10^-4^) or sensory (Δ=−370 ms, *P*=4×10^-3^) cortex in recordings OFF therapy (Supplementary Fig. 3A). Importantly, the significant dopaminergic modulation of motor intention was spatially specific for motor cortex (Δ=−200 ms, *P*=0.029), but not in sensory (Δ=40 ms, *P*=0.81) or parietal (Δ=−7 ms, *P*=0.96) cortex (Supplementary Fig. 2D).

### Dopaminergic medication and STN-DBS shift cortico-subthalamic coupling to theta rhythms

To elucidate neurophysiological correlates of the therapy induced reduction in motor intention to execution, we investigated cortico-subthalamic oscillatory coupling and directionality of information flow with Granger causality. Time-frequency estimates of Granger causality were extracted for motor cortex-STN channel pairs within the motor preparation period (−2 to 0 s). Surrogate estimates from time-reversed signals were subtracted to normalize the resulting time-frequency estimates while reducing the impact of measurement noise.^22^ A previously reported cortico-subthalamic driving of beta oscillations^23^ is visible throughout all conditions with an appreciable but temporally inconsistent difference between OFF therapy and ON levodopa states at end of baseline resting period (−2 s; Figure 3A). At the *time of motor intention*, the ON levodopa condition was associated with a significant cluster of cortico-subthalamic coupling in the theta band (4-10 Hz; peak significance at 7 Hz, −0.9 s, *P*≤0.05 cluster corrected) when compared to the OFF therapy condition (Figure 3C,E). Granger causality at the peak OFF-ON difference correlated with intention to execution delays, as measured as the *time of motor intention*, in the levodopa ON (Spearman’s ρ=0.6, *P*=9×10^-3^; Figure 3G) but not levodopa OFF state (*P*>0.05; not shown). Moreover, comparison of ON STN-DBS with OFF therapy conditions revealed a similar cluster with increased cortico-subthalamic coupling in the theta frequency band (*N*=8; 4-8 Hz; peak significance at 6 Hz, −0.9 s; *P*≤0.05 cluster corrected; Figure 3D,F). Thus, both dopaminergic medication and STN-DBS modulated cortico-subthalamic oscillatory communication with a spectrally specific shift from resting beta to premotor theta coupling, potentially reflective of a shift to a prokinetic circuit state that can accelerate movement initiation in Parkinson’s disease. In summary, this suggests that dopaminergic medication and STN-DBS may share similar mechanisms in the modulation of cortico-subthalamic communication that facilitates action initiation.

**Figure 3.**
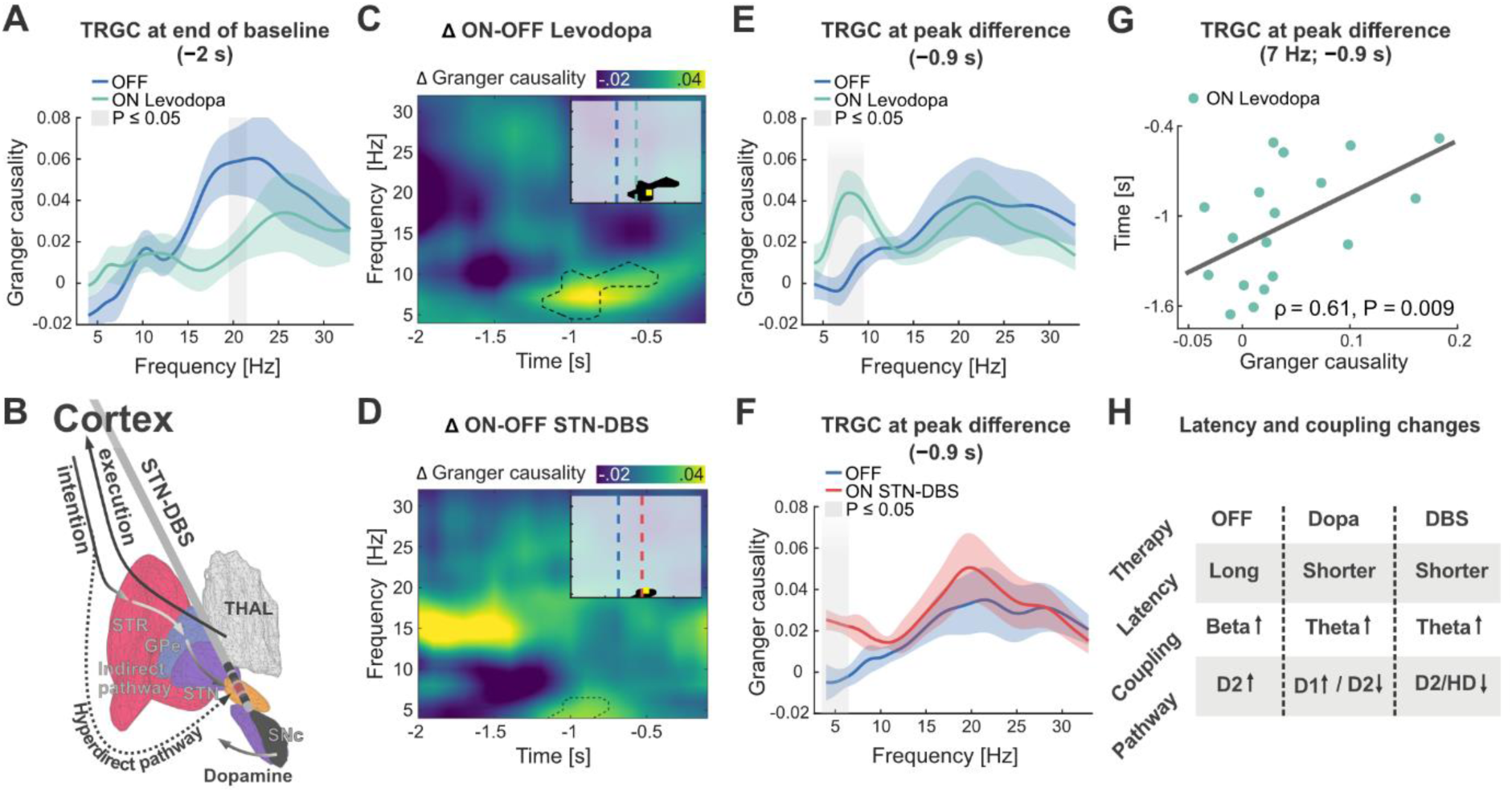
Dopamine and subthalamic deep brain stimulation drive cortico-subthalamic theta coupling that is correlated with shorter intention to execution latencies. **(A)** At the end of the baseline period (-2 s) motor cortex drives oscillatory cortico-subthalamic coupling in the beta frequency range as measured by time-reverse corrected Granger causality (TRGC). **(B)** With start of movement intention motor cortex may drive activity through the trisynaptic striatopallidal indirect and monosynaptic hyperdirect pathways to the subthalamic nucleus (STN), that can be modulated by dopaminergic afferences from SNc and subthalamic deep brain stimulation (STN-DBS) THAL: Thalamus. SNc: Substantia nigra pars compacta. GPe: Globus pallidus externus. **(C)** Comparison of Granger causality OFF vs. ON Levodopa reveals stronger coupling in the theta range (dashed cluster, *P*≤0.05). Peak significance occurred 900 ms before movement onset at 7 Hz (yellow rectangle in inlet), shortly after average decoding time onset (vertical dashed line in blue/green for OFF/ON in inlet). **(D)** A similar effect was observed when comparing OFF therapy cohorts with the sub-cohort ON STN-DBS with a peak significance 900 ms before movement at 6 Hz. Granger causality spectra demonstrate spectrally specific differences for dopamine **(E)** and STN-DBS **(F)** in the theta frequency range. **(G)** Granger causality at peak difference correlated with the *time of motor intention* in the ON Levodopa, but not OFF therapy cohorts. **(H)** In the hypodopaminergic Parkinson’s disease state, intention to execution latency is prolonged and cortico-subthalamic coupling is predominantly observed in the beta frequency range, that could be attributed to excessive indirect pathway (D2) activity. Dopamine and STN-DBS shorten motor intention to execution latencies and shift cortico-subthalamic coupling to the theta frequency band, potentially through suppression and consecutive changes in direct to indirect pathway balance. HD: Hyperdirect pathway.

## Discussion

Our study demonstrates that dopamine and subthalamic deep brain stimulation (STN-DBS) can accelerate and shift the neural dynamics for volitional action in patients with Parkinson’s disease. We found that both motor cortical readiness potentials and oscillatory activity that precede self-initiated movement execution show long latencies before EMG onset in the hypodopaminergic OFF state. These latencies could be shortened with dopamine and STN-DBS therapy. This provides evidence that cortical motor representations, including readiness potentials, are modulated by dopamine and STN-DBS. When analysing cortico-subthalamic oscillatory coupling, we found that both dopamine and STN-DBS could shift cortico-subthalamic dynamic oscillatory drive in the motor preparation phase from beta to theta frequencies. These findings could link the modulation of neural dynamics of motor preparation to shared circuit mechanisms of dopamine and STN-DBS for the alleviation of impaired action initiation in Parkinson’s disease.

### Human evidence for a role of dopamine in volitional action initiation

After movement related activity of dopaminergic neurons was first described for movement execution, it was later often conceptualized to result from a subjective encoding of sensory input, that was formalized as the well-known reward prediction error.^24,25^ Since then, dopamine research has often been focused on reward processing. Nevertheless, the investigation of movement related dopamine release has recently experienced a renaissance, primarily through methodological advances in animal research.^26^ In the last decade multiple impactful studies have clearly demonstrated the phasic increase of firing and release of dopamine prior to action initiation that was unrelated to sensory stimuli or reward contingency.^5,27^ Importantly, optogenetic and spontaneous activation of dopaminergic neurons has repeatedly been shown to facilitate the neural process underlying movement initiation in rodents.^27^ While direct cell-type specific recordings of dopaminergic activity remain unavailable in human subjects, the unique access to invasive brain signals in patients with Parkinson’s disease undergoing neurosurgery for deep brain stimulation has allowed us to characterize the effects of dopamine on the neural representation of motor preparation signals. Our findings corroborate a role for dopamine in the modulation of motor intention coding and provide direct evidence for the involvement of subcortical brain circuits in the generation of cortical readiness potentials and oscillatory correlates of motor intention in humans. Dopamine decreased the time between onset of cortical readiness potentials and oscillatory activity related to motor intention and the onset of motor execution, indicative that cortex–basal ganglia circuits are involved in their processing. Of note, while dopamine may have additional unspecific effects directly on cortex, STN-DBS could reproduce this effect and is a causal targeted subcortical intervention. Most recently, it has been proposed that action related dopamine signals may relate to neural reinforcement of action, putatively through the calculation of action prediction errors.^28,29^ Dopamine and subthalamic DBS may thus reinstate neural reinforcement for action initiation, vigour generation and neural learning.

### Increased intention to execution latency may underlie akinesia in Parkinson’s disease

Akinesia, the impairment to initiate movement, is a pathognomonic clinical feature of the hypodopaminergic parkinsonian state. Investigation of akinesia in experimental settings has so far been largely limited to reaction time experiments that included one or multiple alternative sensory cues.^30^ Both dopaminergic medication and STN-DBS can shorten reaction times, but effects were reported to vary with complexity of the tasks that were primarily defined by sensory input.^30^ Meanwhile, it is known that visual cues can help patients to overcome akinesia, which may indicate that movement initiation in response to sensory cues is less affected than self-initiation in Parkinson’s disease. Experimental studies have provided some evidence for this,^31–33^ but in self-initiated tasks often only surrogate behavioural markers after movement onset were measured as it is inherently difficult to develop a behavioural predictor of the intention to move in a fully self-initiated movement. Therefore, some studies have instead measured electroencephalographic readiness potentials for the investigation of akinesia, mostly focusing on signal magnitudes that were initially reported to be increased in Parkinson’s disease but followed by conflicting results.^31,34–41^ Similar to our approach, one study used EEG during self-paced movements and found longer latencies between readiness potential onset in Parkinson’s disease patients compared to healthy controls.^35^ This suggests that motor preparation in cortex takes longer in the parkinsonian state, which is further supported by reports of prolonged premotor excitability in Parkinson’s disease patients compared to healthy controls, ^42^ and in studies of macaques with neurotoxin-induced parkinsonism that found evidence for longer delays between onset of motor cortex activity and movement.^43,44^ Our study significantly extends these findings by using invasive cortical recordings combined with brain signal decoding to demonstrate that both readiness potentials and oscillatory representations preceding self-initiated movement can be shortened by dopamine and STN-DBS in Parkinson’s disease and providing a neural circuit mechanism of this these effects through modulation of oscillatory cortex – basal ganglia communication with therapy induced increases in cortical theta drive to the subthalamic nucleus.

### Dopamine and deep brain stimulation shift cortico-subthalamic circuit communication to prokinetic oscillatory states

The inhibitory akinetic state in Parkinson’s disease has previously been linked to excessive indirect pathway activity that is correlated with pathological beta band oscillations (13-35 Hz) in Parkinson’s disease. Beta oscillations and their coupling have been proposed to signal the maintenance of a “status quo” that is suppressed by behavioural, motor and cognitive state changes.^45^ Importantly, cortex drives beta activity in the basal ganglia, which can develop a vulnerability to beta hypersynchrony in the absence of dopamine.^23^ An impactful rodent study has demonstrated this vulnerability in individually identified striatal medium spiny neurons in vivo in a dopaminergic lesion model of Parkinson’s disease.^46^ Compared to healthy animals, striatal indirect pathway medium spiny neurons of animals with dopaminergic lesions showed increased firing rates and excessive phase locking to cortical beta oscillations. Another study found that in healthy rodents, strong theta oscillations are present in striatum (∼8 Hz) and associated with movement.^47^ In our study, we demonstrate that dopamine and STN-DBS can shift oscillatory cortico-subthalamic drive from beta to theta rhythms. In the context of perceptual decision making, cortico-subthalamic theta activity has been associated with the modulation of decision thresholds in the presence of conflict.^48^ Thus, one could speculate that similar threshold mechanisms govern the intention to execution period of simple self-initiated movements that could relate to a balance of direct (theta) and indirect pathway (beta) activation. A more recent study has found that cortical beta oscillations could track perceived effort, while theta oscillations related to previous rewards during decision making.^49^ The observed shift to cortico-subthalamic theta drive related to therapeutic alleviation of akinesia in our study could thus be conceptualized as an overcoming of pathologically exaggerated effort perception and decreased reward prediction at the circuit level, which could be rebalanced with dopamine and DBS to accelerate motor initiation. While this remains speculative, it provides an intriguing conceptual foundation to interpret oscillatory changes in hyperkinetic states, where excessive theta and low frequency activity in the basal ganglia has been associated with tics in Tourette’s syndrome and involuntary muscle contraction in dystonia.^9^ Similarly, previous human studies in Parkinson’s disease patients have shown that a shift from an akinetic to a dyskinetic state with involuntary movements as a side-effect of levodopa can be accompanied by a persistent change in oscillatory patterns from beta to theta as the dominant rhythm of the basal ganglia,^50,51^ but these studies mostly investigated patient states rather than dynamic modulations. Notably, one study has reported that phasic theta increases in the subthalamic nucleus during movement may be associated with decreased symptom severity in the parkinsonian ON state, further support for our observation that theta synchrony can be associated with shorter intention to execution latencies.^52^ Taken together, our results corroborate a role for theta oscillations in the reduction of intention to execution delays and their underlying neural processes. While beta oscillations may signal the maintenance of a “status quo” or increased effort perception in the absence of dopamine, theta oscillations may signal the deviation of a “status quo” that can be associated with increased reward predictions during dopamine release that can be facilitated by deep brain stimulation.

### Motor intention decoding for the future of adaptive DBS

This study demonstrates for the first time that it is possible to accurately predict the intention to move from electrocorticography (ECoG) activity in Parkinson’s disease patients performing self-initiated movements. This may have significant implications on the development of adaptive DBS systems, in which stimulation parameters are adapted to the patient’s behavioural state or to neural physiomarkers in real-time. First multicentre trials have been initiated to test the clinical utility of subthalamic beta power as a feature for adaptive stimulation control algorithms. Additionally, it has been proposed that machine learning based symptom decoding using multivariate features from cortical or subthalamic electrodes may extend this approach,^53–56^ especially because beta activity is suppressed by both voluntary and involuntary movements, such as tremor, in Parkinson’s disease. Our study showcases high prediction accuracies of motor intention in cortical but not subthalamic signals. In the future, decoding of movement intention could inform adaptive stimulation algorithms to support movement initiation. Given that DBS suppresses STN activity and thus the indirect pathway,^57^ DBS may have similar effects to a phasic dopamine release. We might speculate that this could not only improve movement intention but reenact the connection of volition and neural reinforcement that is lost in the parkinsonian OFF state. Through this, motor intention-based adaptive DBS could represent a neuroprosthetic support of dopamine circuits for invigoration and reinforcement of volitional action. If successful, it could usher a paradigm shift away from the demand-dependent suppression of “noisy signals”,^58^ towards the targeted restoration of cortex-basal ganglia communication, precisely when it is required.

## Conclusion

In conclusion, we provide direct human evidence that dopamine and the basal ganglia can modulate preparatory brain signals for the self-initiation of volitional action in people with Parkinson’s disease. Our findings suggest that the hypodopaminergic parkinsonian OFF state is associated with increased latencies between the earliest cortical representations of movement initiation and the onset of motor execution. Dopamine and subthalamic deep brain stimulation shortened these latencies which could underlie their effectiveness in alleviating the impairment to initiate movements in Parkinson’s disease patients. This shortening was accompanied by a shift of neural dynamics and brain network connectivity, from antikinetic beta towards prokinetic theta coupling between motor cortex and subthalamic nucleus. Our study provides mechanistic insights into the interplay of dopamine and cortex–basal ganglia communication in the sequence from motor intention to movement execution. In the future, our study could inspire a clinical paradigm shift, away from the suppression of basal ganglia noise, towards targeted network interventions that support intrinsic capacity of dopaminergic circuits for invigoration and reinforcement of purposeful action.

## Data availability

Data from Berlin can be made available conditionally to data sharing agreements in accordance with data privacy statements signed by the patients within the legal framework of the General Data Protection Regulation of the European Union.

## Acknowledgements

We thank all patients who participated in this study. Without their dedication to contribute to the understanding and treatment of Parkinson’s disease, our research would not be possible.

Computation has been performed on the HPC for Research cluster of the Berlin Institute of Health.

## Funding

The study was funded by Deutsche Forschungsgemeinschaft (DFG, German Research Foundation, Project-ID 424778371 – TRR295). W.J.N. received funding from the European Union (ERC, ReinforceBG, Project 101077060). A.A.K. has received grant funding from the Lundbeck foundation as part of the collaborative project grant "Adaptive and precise targeting of cortex-basal ganglia circuits in Parkinsońs disease” (Grant Nr. R336-2020-1035). The NeuroCure Research Centre is Funded by the Deutsche Forschungsgemeinschaft (DFG, German Research Foundation) under Germanýs Excellence Strategy (EXC-2049 – 390688087).

## Competing interests

A.A.K. reports personal fees from Medtronic and Boston Scientific. G.H.S. reports personal fees from Medtronic, Boston Scientific, and Abbott. W.J.N. serves as consultant to InBrain and reports personal fees from Medtronic.

## Supplementary material

Supplementary material is available at *Brain* online.

## Notes

https://github.com/richardkoehler/paper-motor-intention

### Summary of Updates

Minor changes to figures The abstract has been rewritten. Minor corrections in the main text.

