## Supplements for "Dopamine and DBS accelerate the neural dynamics of volitional action in Parkinson’s disease"

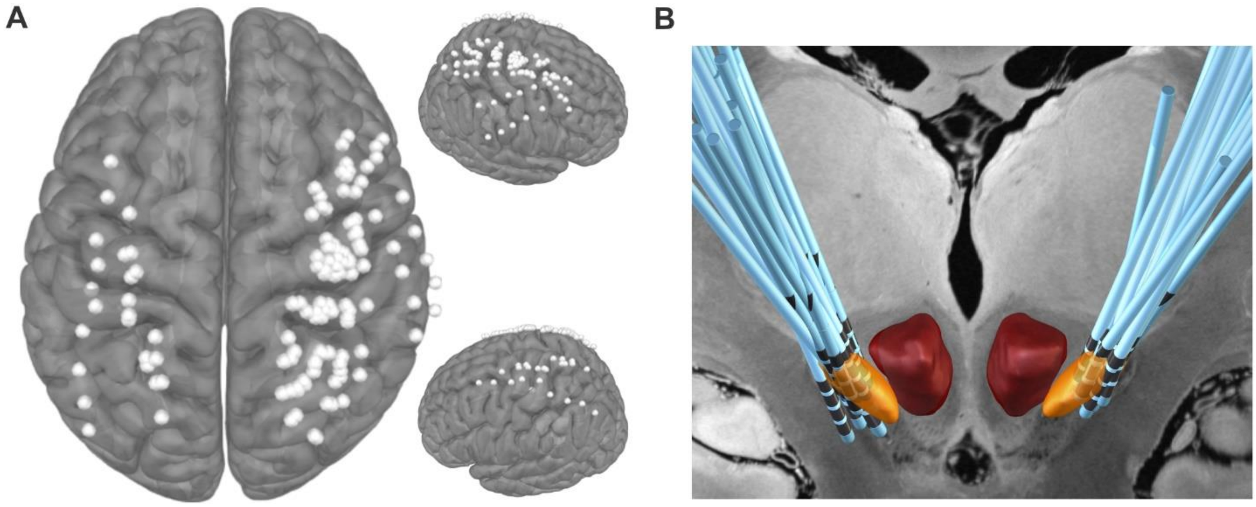


**Supplementary Figure 1 Localisation of electrocorticography and deep brain stimulation electrodes. (A)** Bipolar derivations of all electrocorticography electrodes, and **(B)** deep brain stimulation electrodes after localization and normalization to the MNI (Montreal Neurological Institute) ICBM 2009c Nonlinear Asymmetric template. Orange: subthalamic nucleus, red: Red nucleus.

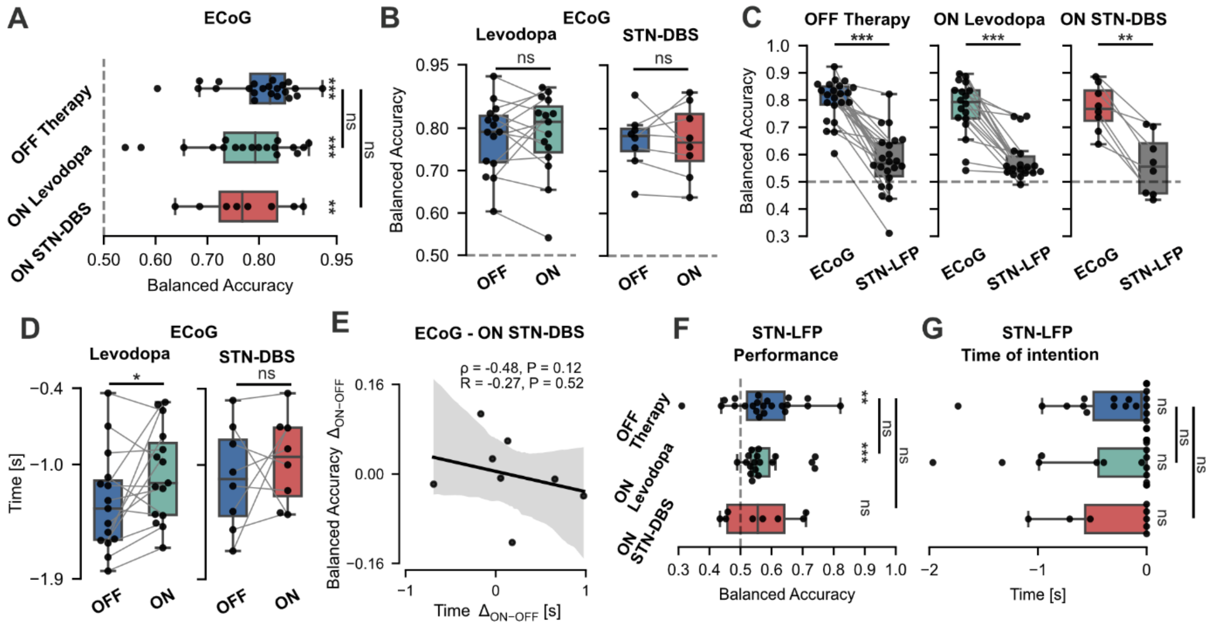


**Supplementary Figure 2 Supplementary analyses for motor intention decoding performance and time of motor intention.** **(A)** Classification performance of single-subject electrocorticography (ECoG) classifiers of all subjects and **(B)** only subjects with repeated recordings. **(C)** Comparison of classification performance of ECoG vs subthalamic local field potential (STN-LFP) based classifiers. **(D)** Time of motor intention derived from ECoG classifier outputs of subjects with repeated recordings. **(E)** Classification performance versus time of motor intention, plotted as difference ON minus OFF, for subjects with recordings both OFF therapy and ON subthalamic deep brain stimulation (STN-DBS). Spearman's $\rho$ and Pearson's $R$ displayed. **(F)** Classification performance and **(G)** time of motor intention of single-subject STN-LFP classifiers. $*P\leq0.05$; $**P\leq0.01$; $***P\leq0.001$

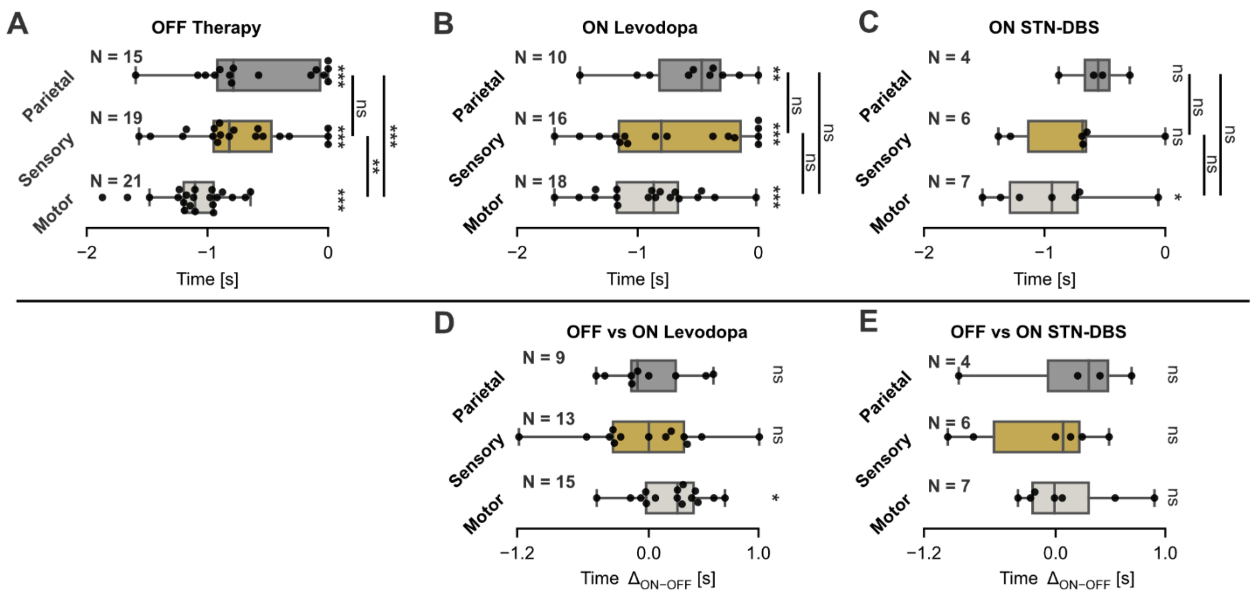


**Supplementary Figure 3 Region-wise comparison of time of motor intention. (A)** Region-wise comparison of time of motor intention OFF therapy, **(B)** ON levodopa, and **(C)** ON subthalamic deep brain stimulation (STN-DBS**). (D)** Region-wise difference of time of motor intention in subjects recorded both OFF and ON levodopa, and **(E)** both OFF and ON STN-DBS. **P*≤0.05; ***P*≤0.01; ****P*≤0.001

**Supplementary Table 1: Clinical scores during recording sessions**

| ID | Session UPDRS-III Med. OFF | Session UPDRS-III Med. ON |
|---|---|---|
| EL003 | 20 | 15 |
| EL004 | 32 | 18 |
| EL005 | 20 | 10 |
| EL006 | 27 | 15 |
| EL007 | 31 | 22 |
| EL008 | 25 | 18 |
| EL009 | 15 | 7 |
| EL010 | 30 | 27 |
| EL011 | 43 | 33 |
| EL012 | 27 | 9 |
| EL013 | 27 | 9 |
| EL014 | 35 | 13 |
| EL015 | 47 | 28 |
| EL016 | 23 | 14 |
| EL017 | 37 | 34 |

UPDRS-III: Unified Parkinson's Disease Rating Scale Part III; Med.: dopaminergic medication

**Supplementary Table 2: Stimulation parameters and clinical scores at 12-month follow-up**

| ID | UPDRS-III 12MFU DBS OFF | UPDRS-III 12MFU DBS ON | DBS contacts | DBS amplitude [mA] |
|---|---|---|---|---|
| EL002 | 57 | 30 | L: 1+8-<br>R: None | L: 0.8 R: None |
| EL005 | 20 | 13 | L: 2+3+4+<br>R: 5+6+7+ | L: 2.0 R: 3.0 |
| EL006 | 57 | 24 | L: None<br>R: 2+3+4+ | L: None R: 2.0 |
| EL007 | 48 | 35 | L: 2+3+4+<br>R: 2+3+4+ | L: 2.0 R: 1.5 |
| EL008 | 25 | 16 | L: 7+8+9+<br>R: 7+8+9+ | L: 2.0 R: 2.0 |
| EL009 | 35 | 11 | L: 2+3+4+<br>R: 2+3+4+ | L: 2.0 R: 2.0 |
| EL012 | 43 | 27 | L: 2+3+4+<br>R: 5+6+7+ | L: 1.5 R: 1.5 |
| EL017 | n/a | n/a | L: 2+3+4+<br>R: 2+3+4+ | L: 3.0 R: 2.0 |

UPDRS-III: Unified Parkinson’s Disease Rating Scale Part III; DBS: deep brain stimulation; L: left; R: right; 12MFU: 12-month follow-up; n/a: not available (12-month follow-up not yet taken place at time of manuscript submission)
